# Multimodal principal component analysis to identify major features of white matter structure and links to reading

**DOI:** 10.1101/2020.05.04.076521

**Authors:** Bryce L. Geeraert, Maxime Chamberland, R. Marc Lebel, Catherine Lebel

**Affiliations:** Biomedical Engineering Graduate Program, University of Calgary; Alberta Children’s Hospital Research Institute, University of Calgary; Cardiff University Brain Research Imaging Centre (CUBRIC), School of Psychology, Cardiff, United Kingdom; Department of Radiology, University of Calgary; GE Healthcare

## Abstract

The role of white matter fibers in reading has been established by diffusion tensor imaging (DTI), but DTI cannot identify specific microstructural features driving these relationships. Neurite orientation dispersion and density imaging (NODDI), inhomogeneous magnetization transfer (ihMT) and multicomponent driven equilibrium single-pulse observation of T1/T2 (mcDESPOT) can be used to link more specific aspects of white matter microstructure and reading due to their sensitivity to axonal packing and fiber coherence (NODDI) and myelin (ihMT and mcDESPOT). We applied principal component analysis (PCA) to combine DTI, NODDI, ihMT and mcDESPOT measures (10 in total), identify major features of white matter structure, and link these features to both reading and age. Analysis was performed for nine reading-related tracts in 46 neurotypical 6-16 year olds. We identified three principal components (PCs) which explained 79.5% of variance in our dataset. PC1 probed tissue complexity, PC2 described myelin and axonal packing, while PC3 was related to axonal diameter. Mixed effects models regressions did not identify any relationships between principal components and reading skill. Further Bayes factor analysis revealed that absence of relationships was not due to low power. PC1 suggested increases in tissue complexity with age in the left arcuate fasciculus, while PC2 suggested increases in myelin and axonal packing with age in the bilateral arcuate, inferior longitudinal, inferior fronto-occipital fasciculi, and splenium. Multimodal white matter imaging and PCA produce microstructurally informative, powerful principal components which can be used by future studies of development and cognition.

## 1 INTRODUCTION

Reading is a sophisticated skill with many constituent systems including vision, language, memory, and attention. White matter fibers play an important role in connecting these systems and facilitating coordinated processing across the reading network. Diffusion tensor imaging (DTI) is frequently used to investigate links between white matter and reading thanks to its sensitivity to white matter microstructural features. DTI studies have linked white matter to reading in a broad network of tracts including the arcuate, superior and inferior longitudinal, inferior fronto-occipital, and uncinate fasciculi, and the posterior corpus callosum [1–5], such that markers of increased white matter maturity correlate with improved reading scores. Additionally, longitudinal DTI studies show that maturation of reading-related tracts is related to improvements in reading ability [6–10]. White matter abnormalities have been observed in children with reading difficulties, most often in left temporo-parietal white matter [11–14] as language and reading networks are typically left lateralized [11, 15, 16]. Finally, changes in DTI measures are observed in reading-related white matter following reading interventions [17–19].

DTI studies have identified a network of white matter related to reading but cannot comment on the particular features of white matter microstructure driving these relationships. Fractional anisotropy (FA) and mean diffusivity (MD) describe total water diffusion and are simultaneously sensitive to many microstructural factors [20–23]. Newer techniques with increased specificity may be used to build upon DTI literature. Neurite orientation dispersion and density imaging (NODDI) produces the neurite density index (NDI) and orientation dispersion index (ODI) which are sensitive to axonal packing and tract coherence, respectively [24]. Inhomogeneous magnetization transfer (ihMT) and multicomponent driven equilibrium single-pulse observation of T1 and T2 (mcDESPOT) produce the quantitative ihMT (qihMT) and myelin volume fraction (VF_m_) measures respectively, both sensitive to myelin [25, 26]. Additionally, measures of axon volume and myelin volume such as NDI and VF_m_ can be combined to produce the g-ratio, which describes the ratio of axon thickness to total fiber diameter [27]. These methods have been validated in *in vitro* studies [28–33], and they hold great potential to clarify our understanding of white matter development and links to reading.

Investigating multiple imaging measures in a univariate fashion, the typical practice in developmental studies to date, necessarily increases the required stringency of multiple comparisons corrections, thereby reducing the discriminating power of the analysis. One solution to preserve power and reduce comparisons is to collapse white matter measures into orthogonal components via principal component analysis (PCA). A framework using PCA for dimensionality reduction in white matter has been recently described [34], and resultant components were linked to age, suggesting developmental sensitivity. The goal of this study was to combine white matter imaging techniques (DTI, NODDI, ihMT, and mcDESPOT) to better understand relationships between brain structure and reading in a sample of healthy 6-16 year old children. Based upon the PCA results of Chamberland et al., we hypothesized that observed principal components would represent diffusion restriction and tissue complexity factors, and that these components would be linked to reading proficiency in reading-related tracts, such that indications of more myelin and axonal packing would be related to better reading performance.

## 2 METHODS

### 2.1 Participants

46 healthy participants aged 6-16 years (mean age: 11.0 ± 2.6 years, 24 males / 22 females) were recruited as part of an ongoing study on adolescent brain development. Inclusion criteria were: 1) uncomplicated birth between 37-42 weeks’ gestation, 2) no history of developmental disorder, psychiatric disease, or reading difficulty, 3) no history of neurosurgery, and 4) no contraindications to MRI. 22 children (mean age: 13.3 ± 2.6 years, 11 males / 11 females) returned 2 years after their initial visit for a second scan and cognitive assessment. All subjects provided informed assent and parents/guardians provided written informed consent. Gender was determined by parent report. This study was approved by the local research ethics board (ethics ID: REB13-1346).

### 2.2 Imaging

Subjects were scanned using a 32-channel head coil on a GE 3T Discovery MR750w (GE, Milwaukee, WI) system at the Alberta Children’s Hospital. Two diffusion-weighted datasets were acquired at b = 900 s/mm^2^ and 2000 s/mm^2^ using a spin-echo echo planar imaging sequence with TR/TE = 12s/88ms, 2.2 mm × 2.2 mm × 2.2 mm resolution, with 5 b = 0 s/mm^2^ volumes and 30 gradient directions per volume, scan duration = 14:24 minutes. IhMT images used a 3D SPGR sequence: TR/TE = 10.46ms/2.18ms, 2.2mm × 2.2 mm × 2.2 mm resolution, flip angle 8°, scan time 5:12 minutes. The sequence included a 5ms Fermi pulse with peak B1 of 45 mG and 5kHz offset prior to each excitation. The MT condition cycled between positive offset (+5kHz), dual offset (±5kHz), negative offset (−5kHz), and dual offset. A 32° flip angle reference image with no MT pulse was acquired for quantification. For mcDESPOT, multi-flip angle 3D SPGR images (α = 3°, 4°, 5°, 6°, 7°, 9°, 13°, and 18°) were collected with TR/TE = 9.1ms/3.9ms, 1.7mm × 0.86mm × 1.7mm resolution; IR-SPGR images were collected to correct for B_1_ inhomogeneity using 5° α, TR/TE = 9.1ms/3.9ms, 2.29mm × 0.86mm × 3.4mm resolution; two multi-flip angle bSSFP images were collected at phase 0° and 180° to correct for B_0_ inhomogeneity, with α = 10°, 13°, 16°, 20°, 23°, 30°, 43°, and 60°, TR/TE = 6.6ms/3.2ms, 1.7mm × 0.86mm × 1.7mm resolution. Total scan time for mcDESPOT was 16:35 minutes. T1-weighted anatomical images were also acquired, with TI = 600ms, TR/TE = 8.2ms/3.2ms, 0.8 mm × 0.8 mm × 0.8 mm resolution, scan duration 5:38.

### 2.3 Image Processing

All images were visually inspected for quality assessment and processed separately using appropriate tools before being combined for principal component analysis. T1 images were processed through FreeSurfer 5.3 (http://surfer.nmr.mgh.harvard.edu/) for intensity normalization and brain extraction. ExploreDTI [35] was used for all DTI processing and analysis, including preprocessing for signal drift correction [36], brain extraction, eddy current and motion corrections [37, 38], and registration to skull-stripped T1 images to correct geometric distortions induced by echo-planar imaging. The REKINDLE model was used to produce FA, MD, radial diffusivity (RD), and axial diffusivity (AD) maps for each subject using the b = 900 s/mm^2^ shell only [39]. Whole brain tractography was performed using constrained spherical deconvolution [40] with Lmax = 6, 2mm isotropic seed voxels, 1mm step size, 30 maximum angle of deviation and an acceptable streamline range of 50 to 500mm. Next, semiautomated tractography [41] was performed to segment the arcuate, inferior longitudinal (ILF), inferior fronto-occipital (IFOF), and uncinate fasciculi bilaterally, along with the splenium, as shown in Figure 1. A 11-year old female with high data quality was selected as the exemplar participant for this process; all regions were drawn on this template brain and then registered to other participants’ data for tracking in native space [42]. Processed DTI data was exported to the NODDI Toolbox (http://www.nitrc.org/projects/noddi_toolbox) for calculation of isotropic (f_iso_) and intracellular (f_icvf_, or NDI) volume fractions and ODI.

**Figure 1:**
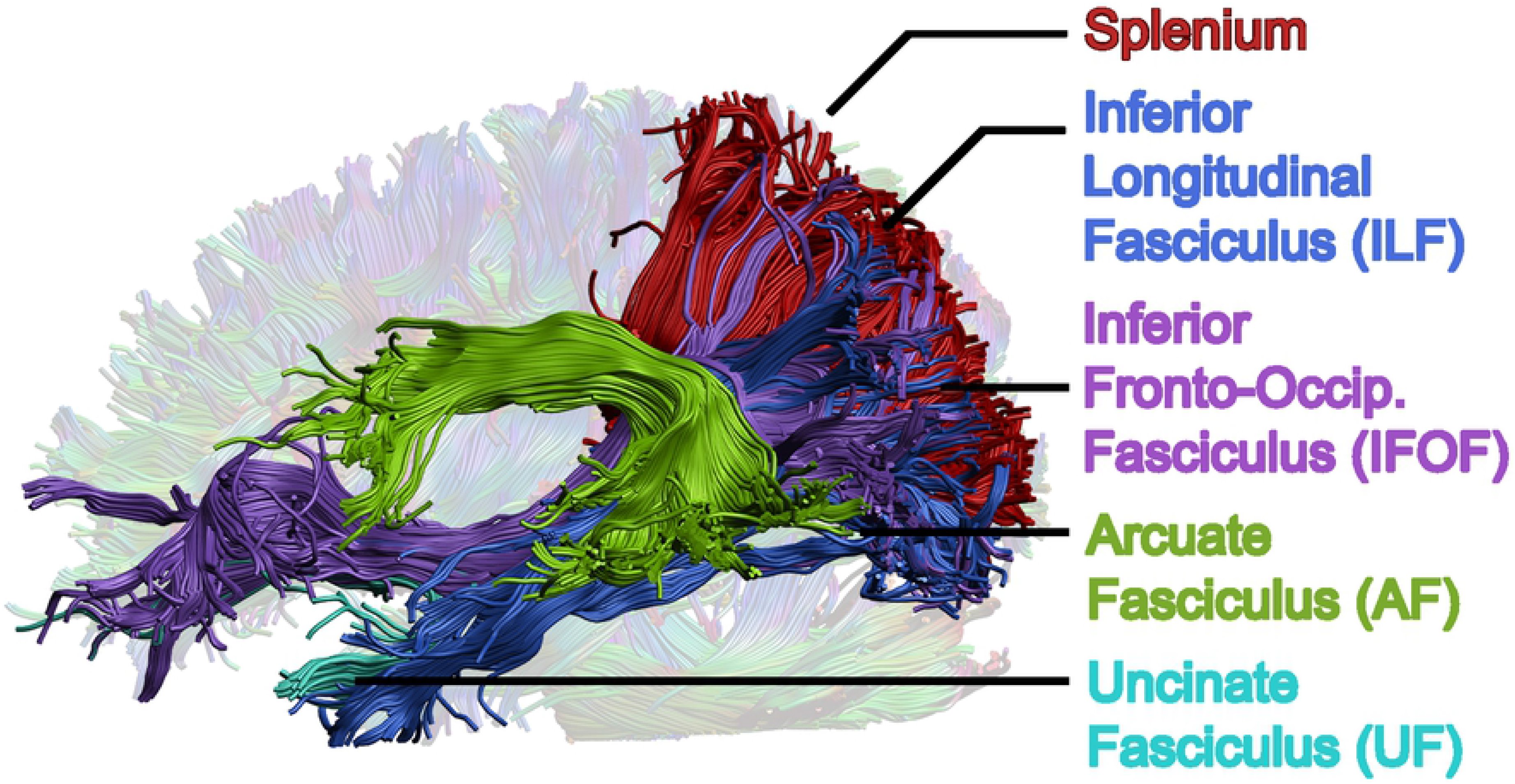
Major, reading-related white matter tracts chosen as regions of interest. Whole brain tractography was performed via constrained spherical deconvolution, then tracts were segmented using deterministic semi-automated tractography in ExploreDTI. Regions of interest were investigated bilaterally, but only the left hemisphere is shown here.

Pseudo-quantitative ihMT maps (qihMT) and magnetization transfer ratio (MTR) maps were obtained from ihMT data using an in-house GE protocol as described in previous work [43]. Following MTR and qihMT image production, brain extraction was performed on MTR images using FSL’s BET2 tool [44], and resulting brain-extracted MTR image was used as a mask to produce a brain-extracted qihMT image.

mcDESPOT SPGR, IR-SPGR, and bSSFP images were aligned to the SPGR image with the largest α then processed by fitting T1, T2, and volume fractions to three water compartments (myelin-bound, intra/extracellular, and free), along with exchange rates between myelin-bound and intra/extracellular water [45]. The myelin-bound water volume fraction from this fitting was used to produce VF_m_ maps for each participant. G-ratio maps were computed using VF_m_, NDI, and f_iso_ maps to calculate the fiber volume fraction (FVF) and g-ratio using the following two equations.

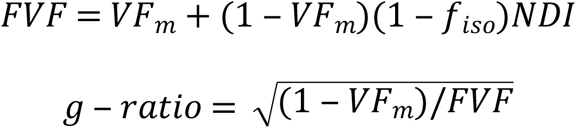

Following production of all measure maps, qihMT, MTR, VF_m_, NDI, and ODI maps were registered to b = 900 s/mm^2^ FA maps using Advanced Normalization Tools (ANTs) [46]. Default parameters from antsRegistrationSyN.sh were used, with the –t s flag chosen to select rigid, affine, and deformable symmetric normalization transforms. Then, the mean FA, MD, AD, RD, NDI, ODI, MTR, qihMT, VF_m_, and g-ratio values were extracted for all 9 segmented tracts per participant. Additionally, along-tract analysis was performed for each tract in ExploreDTI [47, 48], to produce a profile of measure means for all ten measures at twenty equidistant segments per tract.

### 2.4 Reading Assessments

Reading was evaluated using the Wechsler Individual Achievement Test – Third Edition: Canadian [49]. Participants completed the Reading Comprehension, Word Reading, Pseudoword Decoding, and Oral Reading Fluency subtests. From these subtests, the Total Reading Composite Score was computed as a measure of general reading proficiency. This score combines phonological awareness, reading comprehension, and fluency.

### 2.5 Statistical Analysis

All statistical analysis was performed in R version 3.6.1 [50]. Along tract data for each subject’s first time point (10 measures × 9 tracts × 20 segments) was combined into a single table for principal component analysis. The format of this table has been described by Chamberland et al [34]. PCA was performed via the *prcomp* function (using the scale = 1 option to normalize each feature independently). A Kaiser-Meyer Olkin (KMO) test was used to assess sampling adequacy of PCA results [51]. Following PCA, input variable contributions to principal components along with correlations between variables within along-tract data were inspected to identify redundancy between variables. In the case of highly collinear measures (i.e., measures which contributed to PCA outputs in very similar fashions), the variable with highest correlations to all other input measures was removed in order to improve stability of PCA computations [52] and the PCA was recomputed. Resultant principal components with eigenvalue > 1 were retained, while other components were discarded [53]. Varimax rotation was performed on component loadings (the rotations matrix output by *prcomp*) via the *varimax* function to aid in interpretation of principal components. Measures were considered meaningful contributors to a resultant principal component if they accounted for above average variance (>11.1%) in the component.

Following varimax rotation, longitudinal principal component weightings were calculated by multiplication of time point 2 along tract data by the rotation matrix output by *varimax*. Next, along tract weightings for principal components were averaged in each tract to produce mean principal component weightings for each subject in all 9 investigated tracts. Linear mixed effects models were computed via *lmer* to investigate relationships between principal components with Total Reading and age in each tract. Age models included age, gender, an age*gender interaction, and a random intercept per subject. If the age*gender interaction was not significant, it was removed and the model was rerun. Total Reading models for each tract included all retained principal components along with age, and gender if a gender effect was observed for any principal component. Restricted maximum likelihood was used for all models. Benjamini-Hochberg false discovery rate (FDR) correction was used to correct for 27 comparisons (9 tracts × three principal components). Multiple comparisons corrections were conducted separately for age and Total Reading findings. Example formulas are provided below. Time point 1 data for each measure included in our final PCA was correlated with Total Reading via partial correlation in each region, controlling for age, and FDR correction was applied for 9 correlations across each measure.

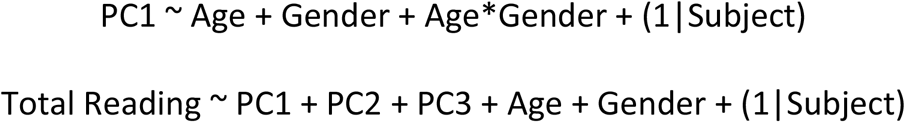

Bayes factor analysis was performed via *generalTestBF* in the BayesFactor package for R [54] to supplement regression analysis by assessing the observed statistical power of models connecting retained principal components and Total Reading. Bayes factors output by *generalTestBF* were inverted to reflect the ratio of likelihood of the null hypothesis divided by the likelihood of a given model. A Bayes factor of greater than 3, indicating our data was 3 times more likely to be described by the null hypothesis than a given model, was considered evidence for the null hypothesis. A Bayes factor of less than 1/3, indicating that a model including our chosen predictors was 3 times more likely to explain our data than the null hypothesis condition, was considered evidence for the alternative hypothesis. Bayes factors between 1/3 and 3 were considered an indication of low power, such that neither evidence for the null or alternative hypotheses could be inferred [55].

## 3 RESULTS

### 3.1 Principal component analysis

Figure 2 visualizes each included imaging metric in the splenium. Here we can see that measures with shared sensitivities vary similarly across the tract. For example, FA, RD, qihMT, and VF_m_ are all similar to myelin and reach extreme values in the center of the splenium (highly positive for FA, qihMT, and VF_m_, highly negative for RD).

**Figure 2:**
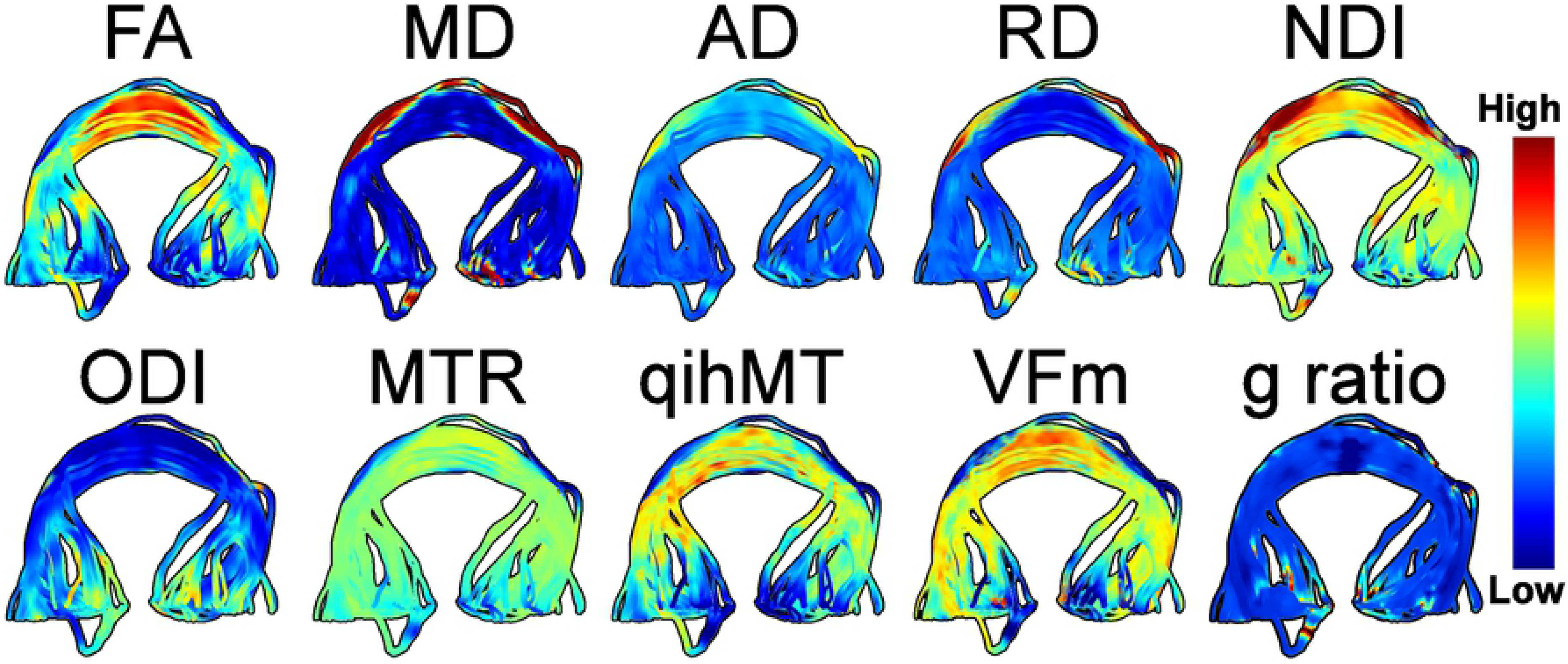
Multimodal imaging of white matter microstructure in the splenium. Measures from DTI, NODDI, MT, and mcDESPOT imaging can be contrasted to provide a multifaceted understanding of white matter structure.

MTR was removed from our principal component analysis due to high collinearity with qihMT (r^2^ = 0.64). Three principal components were identified in our final model, which collectively explained 79.5% of variance (KMO test value = 0.53). Measures contributing greater than 11.1% variance (expected if all variables contributed uniformly) to a component following varimax rotation are visualized in Figure 3. Principal component (PC) 1 explained 37.5% of variance and was primarily composed of measures sensitive to tissue complexity: FA, AD, ODI, along with MD. PC2 explained 23.0% of variance and was composed of measures sensitive to myelin and axon packing: FA, MD, RD, and NDI. PC3 explained 19.0% of variance and was driven by measures sensitive to myelin and axonal diameter, VF_m_ and g-ratio.

**Figure 3:**
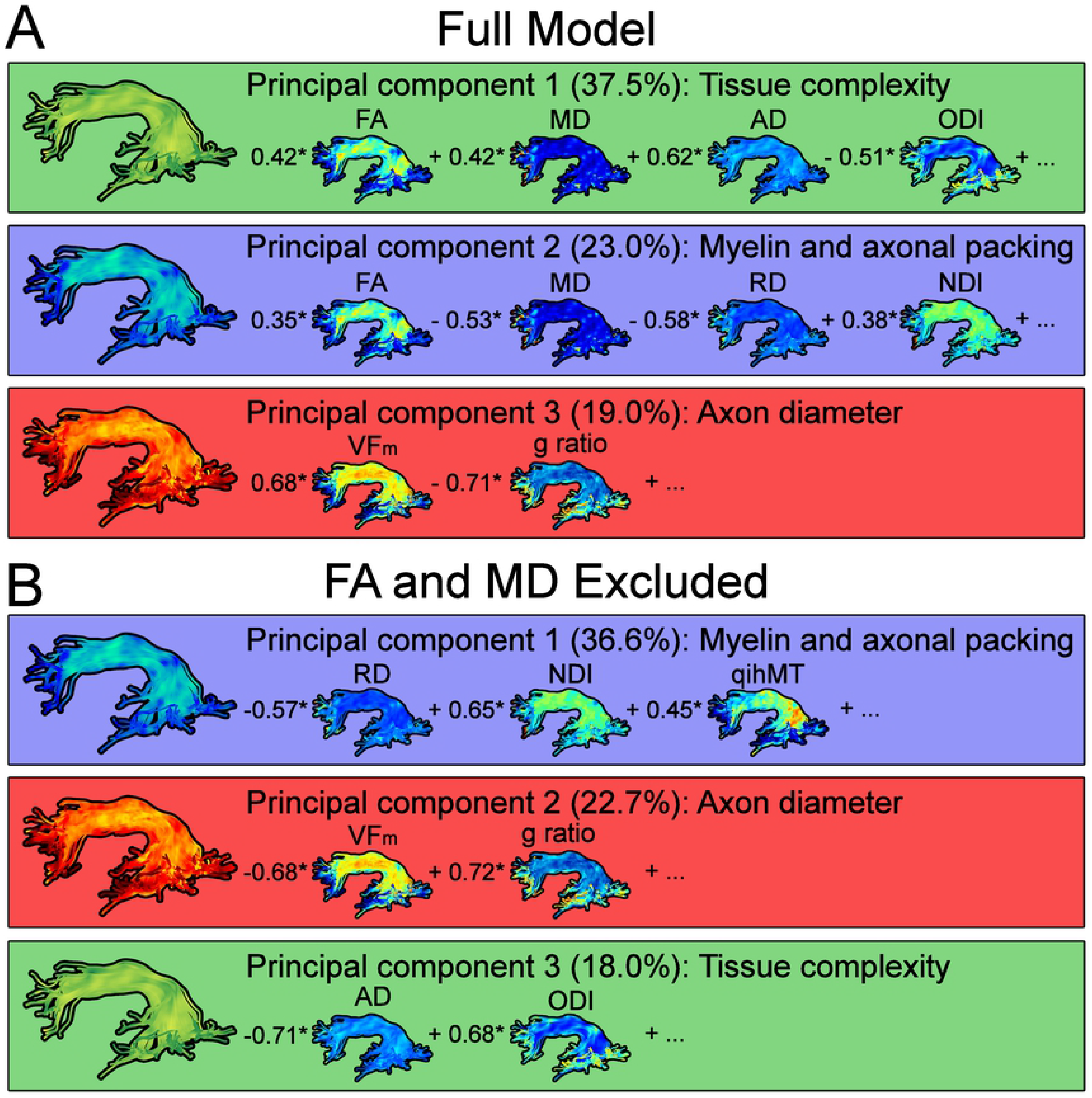
Resultant components from principal component analysis visualized in the left arcuate fasciculus. Correlations for measures which contribute greater variance than expected by chance (>11.1%) are included for each component. Panel A displays PCA results from all 9 measures. Components in Panel A explained 79.5% of variance in our data (variance explained by each individual component is noted in brackets). Principal components were related to diffusion along a primary axis (PC1), myelin and axonal packing (PC2), and axon diameter (PC3). Panel B shows results from a secondary PCA with FA and MD removed. Principal components in Panel B explain 77.3% of variance, and clarify our interpretation of principal components.

As shown in Figure 3 panel A, FA and MD contributed strongly to PC1 and PC2 even after varimax rotation, likely because FA and MD are broadly sensitive to white matter structure. We removed FA and MD and recomputed PCA to interpret our principal components with increased clarity (results shown in Figure 3 panel B). This resulted in a model with three principal components that explained 77.3% of variance (KMO = 0.43), denoted as PC_B_. In this model, PC1_B_ explained 36.6% of variance and was composed of RD, NDI, and qihMT. PC2_B_ explained 22.7% of variance and was composed of VFm and g-ratio. Finally, PC3_B_ explained 18.0% of variance and was driven by AD and ODI. While removing FA and MD and running a reduced PCA model aided in interpretation of our principal components, mixed effects models regressions and Bayes factor analyses were conducted with the full PCA model including FA and MD.

### 3.2 Regression Models

Mixed effects models linking principal components to Total Reading scores are summarized in Table 1. No relationships were observed between principal components and Total Reading. To further investigate the absence of relationships between principal components and Total Reading, we followed up by running mixed effects models between principal components and subtest scores for Reading Comprehension, Word Reading, Pseudoword Decoding, and Oral Reading Fluency. No significant relationships were observed between principal components and reading subtest scores. Correlations between the initial measure set and Total Reading are summarized in Supplementary Table 1. No correlations were observed between individual measures and Total Reading scores.

**Table 1.** Parameters for mixed effects models linking principal components to Total Reading (formula: Total Reading ~ PC1 + PC2 + PC3 + Age + (1|Subject)).

| Region | R<sup>2</sup><br>(marginal) | df | Predictor | Estimate ± SE | t | p |
| --- | --- | --- | --- | --- | --- | --- |
| Left arcuate | 0.026 | 64 | PC1 | -1.17 ± 6.61 | -0.18 | 0.860 |
|  |  |  | PC2 | 5.26 ± 3.86 | 1.37 | 0.178 |
|  |  |  | PC3 | 1.52 ± 2.89 | 0.53 | 0.601 |
|  |  |  | Age | -0.00 ± 0.00 | -0.74 | 0.462 |
| Right arcuate | 0.022 | 64 | PC1 | -6.55 ± 5.42 | -1.21 | 0.232 |
|  |  |  | PC2 | -0.13 ± 3.47 | -0.04 | 0.970 |
|  |  |  | PC3 | -4.06E-2 ± 2.47 | -0.02 | 0.987 |
|  |  |  | Age | -8.06E-4 ± 1.72E-3 | -0.47 | 0.641 |
| Left ILF | 0.026 | 64 | PC1 | -5.19 ± 6.44 | -0.81 | 0.424 |
|  |  |  | PC2 | -0.88 ± 4.06 | -0.22 | 0.829 |
|  |  |  | PC3 | -3.34 ± 2.60 | -1.28 | 0.207 |
|  |  |  | Age | 6.64E-5 ± 1.62E-3 | 0.04 | 0.968 |
| Right ILF | 0.034 | 64 | PC1 | 6.06 ± 5.79 | 1.05 | 0.300 |
|  |  |  | PC2 | 3.12 ± 3.87 | 0.81 | 0.423 |
|  |  |  | PC3 | 0.89 ± 2.30 | 0.39 | 0.701 |
|  |  |  | Age | -7.32E-4 ± 1.64E-3 | -0.45 | 0.656 |
|  |  |  | Gender | 1.37 ± 3.74 | 0.37 | 0.716 |
| Left IFOF | 0.052 | 64 | PC1 | 0.75 ± 5.74 | 0.13 | 0.897 |
|  |  |  | PC2 | 8.87 ± 5.01 | 1.77 | 0.081 |
|  |  |  | PC3 | -1.43 ± 2.61 | -0.55 | 0.585 |
|  |  |  | Age | -0.00 ± 0.00 | -0.79 | 0.435 |
| Right IFOF | 0.046 | 64 | PC1 | -1.81 ± 5.94 | -0.30 | 0.762 |
|  |  |  | PC2 | 6.47 ± 3.99 | 1.62 | 0.110 |
|  |  |  | PC3 | 2.08 ± 2.65 | 0.78 | 0.436 |
|  |  |  | Age | -0.00 ± 0.00 | -0.93 | 0.356 |
| Left uncinate | 0.087 | 64 | PC1 | -5.83 ± 5.43 | -1.08 | 0.287 |
|  |  |  | PC2 | 5.72 ± 4.34 | 1.32 | 0.192 |
|  |  |  | PC3 | -3.85 ± 1.94 | -1.99 | 0.053 |
|  |  |  | Age | -3.78E-4 ± 1.59E-3 | -0.24 | 0.813 |
| Right uncinate | 0.008 | 64 | PC1 | 1.16 ± 6.18 | 0.19 | 0.852 |
|  |  |  | PC2 | 2.54 ± 3.38 | 0.75 | 0.455 |
|  |  |  | PC3 | -0.49 ± 2.45 | -0.20 | 0.844 |
|  |  |  | Age | -1.83E-4 ± 1.59E-3 | -0.12 | 0.909 |
| Splenium | 0.035 | 64 | PC1 | 7.28 ± 4.55 | 1.60 | 0.115 |
|  |  |  | PC2 | 2.07 ± 2.50 | 0.83 | 0.410 |
|  |  |  | PC3 | 1.45 ± 2.77 | 0.52 | 0.603 |
|  |  |  | Age | -0.00 ± 0.00 | -0.28 | 0.778 |

Table 2 summarizes models linking principal components to subject age and gender. A significant relationship between PC1 and age was observed in the left arcuate (t = −2.93, p = 0.004). Increases in PC1 with age suggest that FA, MD, and AD increase with age while ODI decreases in the left arcuate, hinting at increased diffusion restrictions and tissue complexity. A similar relationship was observed in the right arcuate but this finding did not survive multiple comparisons corrections. Positive relationships between PC2 and age were observed in the bilateral arcuate (L: t = 3.70, p < 0.001; R: t = 3.66, p < 0.001), inferior longitudinal fasciculus (L: t = 2.75, p = 0.007; R: t = 3.05, p = 0.003), inferior fronto-occipital fasciculus (L: t = 3.21, p = 0.002; R: t = 3.80, p = 0.003), and splenium (t = 2.31, p = 0.024). Increases in PC2 reflect increases in FA and NDI, and decreases in MD and RD, suggesting increased axon packing and myelin with age. The gender main effect (t = −2.01, p = 0.049) and the age*gender interaction were significant for PC3 in the right inferior longitudinal fasciculus, but neither survived multiple comparisons corrections. Scatterplots are provided in Figure 4 to illustrate relationships between PC1, PC2 and age (panels B and C).

**Table 2.**
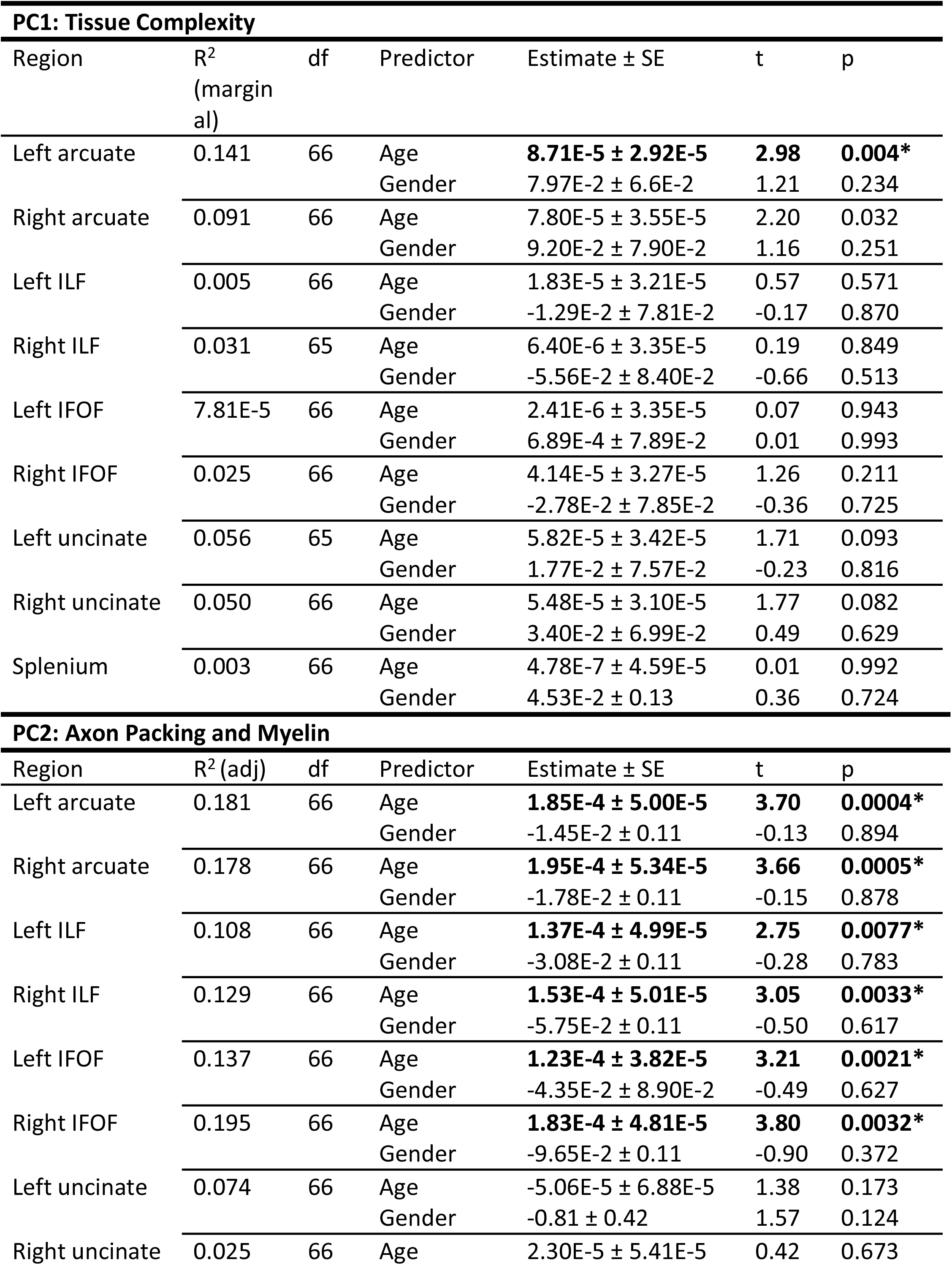
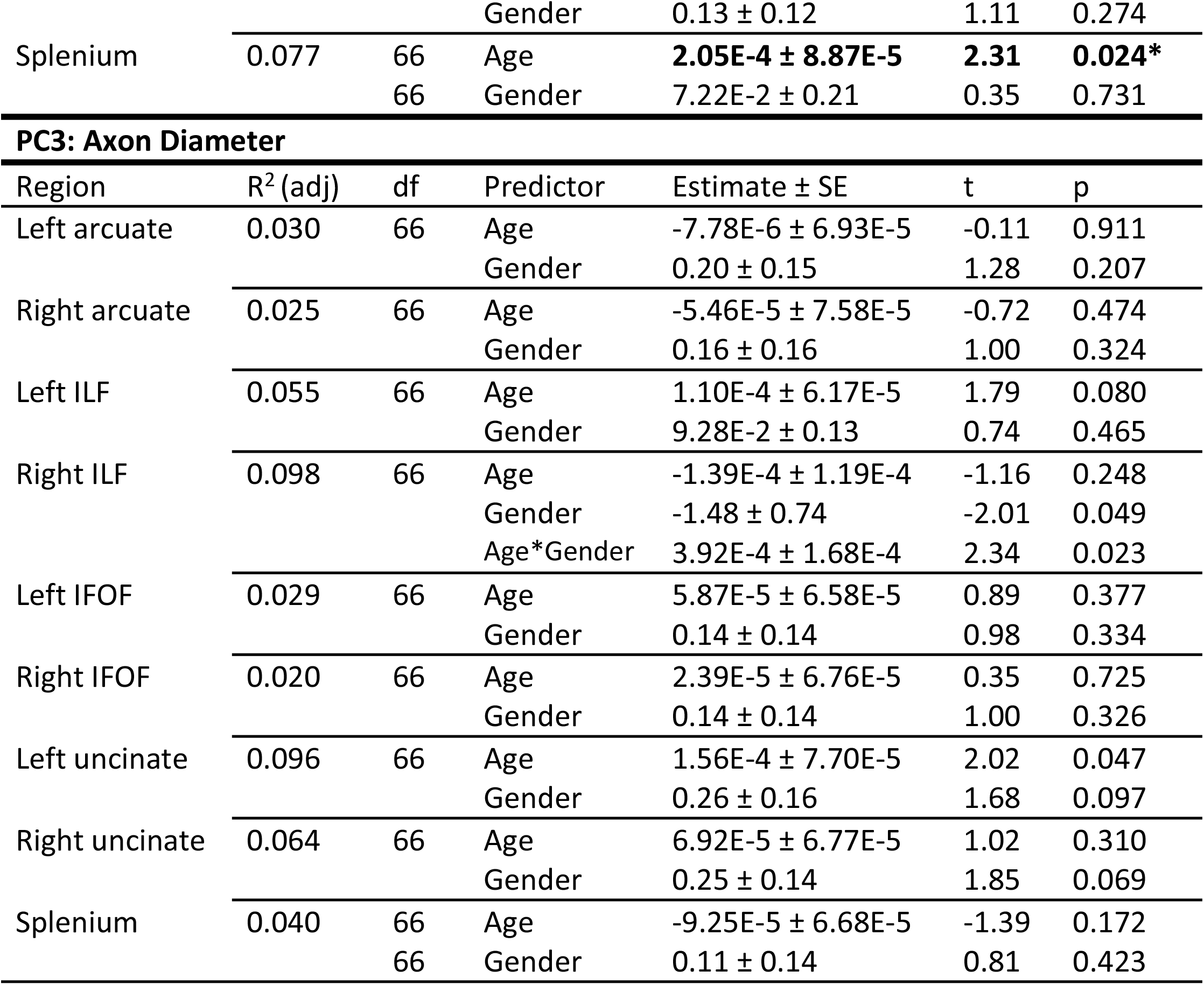
Parameters for mixed effects models regressions linking principal components to age and gender (formula: PC ~ age + gender + age*gender + (1|Subject)). Significant effects that survive multiple comparisons are bolded and marked by an asterisk.

**Figure 4:**
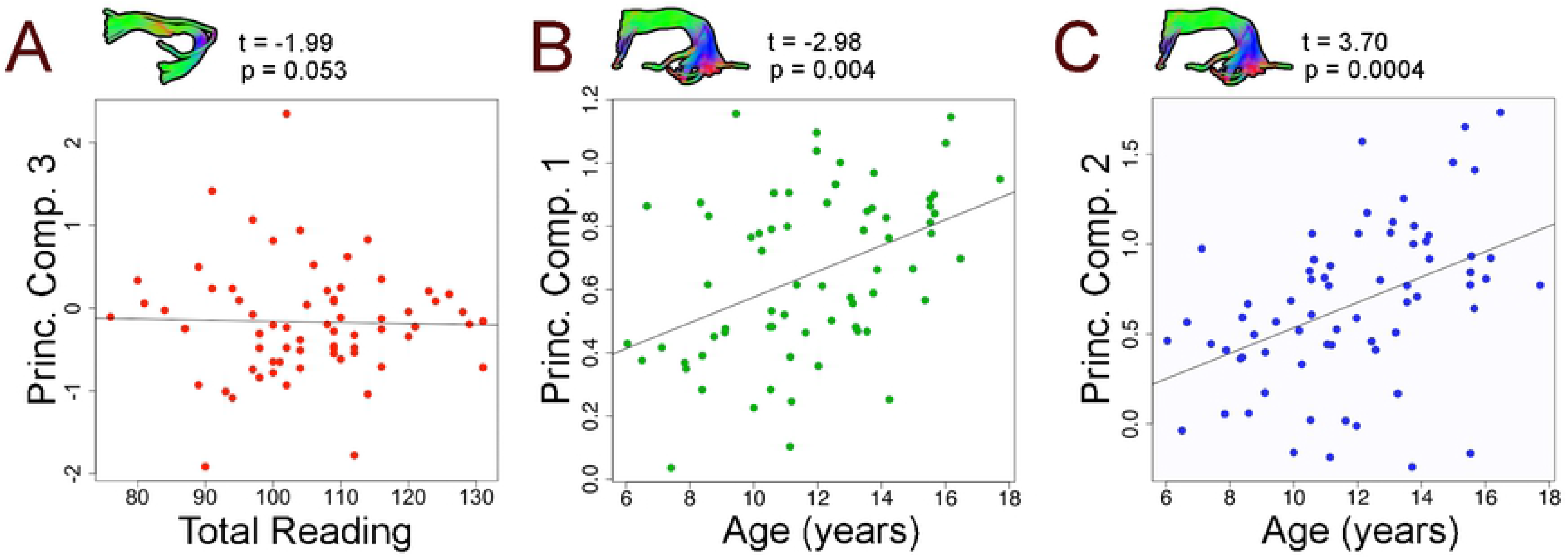
Scatterplots visualizing relationships between principal component 3 (PC3) and Total Reading in the left uncinate fasciculus (A), PC1 and age in the left uncinate (B) and PC2 and age in the left uncinate (C). Increases in PC1 indicate increased diffusion along a primary axis, while increases in PC2 indicate increased myelin and axon packing, thus relationships depicted in panels A and B could potentially reflect axonal maturation. No significant links between principal components and Total Reading were observed. The relationship between PC3 and Total Reading in the left uncinate was closest to our significance threshold.

### 3.3 Bayes Factor Analysis

Bayes factors analysis was conducted to evaluate Total Reading mixed effects models regressions. Results from this analysis are summarized in Table 3. Bayes factors including all principal components and age as covariates of Total Reading were greater than 3 in all regions, indicative of evidence for the null hypothesis.

**Table 3.** Bayes factors assessing the likelihood of the null hypothesis condition (no relationship between Total Reading scores and model components) versus the likelihood of the model condition (relationships between included components and Total Reading). A Bayes factor of 3—indicating our sample data is 3 times more likely to be explained by the null condition than the model condition—or greater provides evidence for the null condition.

| READING MODELS |  |  |
| --- | --- | --- |
| Region | Components | Bayes Factor |
| Left arcuate | PC1 + PC2 + PC3 + Age | 9.43 |
| Right arcuate | PC1 + PC2 + PC3 + Age | 8.93 |
| Left ILF | PC1 + PC2 + PC3 + Age | 47.62 |
| Right ILF | PC1 + PC2 + PC3 + Age | 20.83 |
| Left IFOF | PC1 + PC2 + PC3 + Age | 11.76 |
| Right IFOF | PC1 + PC2 + PC3 + Age | 9.35 |
| Left uncinate | PC1 + PC2 + PC3 + Age | 19.23 |
| Right uncinate | PC1 + PC2 + PC3 + Age | 19.23 |
| Splenium | PC1 + PC2 + PC3 + Age | 10.42 |

## 4 DISCUSSION

Using a multimodal microstructural MR dataset, we identified 3 principal components of white matter structure in reading-related tracts. These principal components represented tissue complexity, axon packing and myelin, and axon diameter. No significant relationships between principal components and Total Reading or components of reading skill were observed. Follow-up Bayes factor analysis provided supplementary evidence for the null hypothesis in all investigated regions. PC1 was negatively linked to age in the left arcuate, and positive relationships between PC2 and age were found throughout the brain. We have shown that multimodal white matter imaging and PCA produce microstructurally informative, powerful principal components which can be used by future studies of development and cognition.

Principal component analysis identified three key components that explained a large proportion of variance (79.5%) in our dataset, and represented tissue complexity (axon coherence), diffusion restriction (axonal packing and myelination), and axon diameter. PC1 explained the largest amount of variance (37.5%). With significant contributions from FA, MD, AD, and ODI, PC1 probed diffusion anisotropy and was most influenced by axon integrity and coherence. PC2 explained 23.0% of variance and reflects myelin and axonal packing, as shown by heavy loadings on FA, MD, RD, and NDI. Finally, PC3 explained 19.0% of variance and is driven by VF_m_ and g-ratio. PC3 likely corresponds to axon diameter as PC2 accounts for a large proportion of variance and contains several myelin-sensitive measures. Studies employing PCA with white matter imaging measures have identified similar principal components related to diffusion anisotropy and overall diffusivity [34, 56]. Our PCA expands upon previous findings by including non-diffusion measures from magnetization transfer and relaxometry. This allowed our multimodal PCA to identify a novel third component related to axon diameter.

Shared information between white matter imaging metrics resulted in multiple principal components loading onto the same measures, in particular FA and MD. This was addressed in multiple ways. First, in the case of highly correlated variables, redundant variables were removed from PCA analysis. Next, varimax rotation minimized loading of multiple principal components onto the same variables, and helped to clearly illustrate differences between resultant principal components. Finally, re-running PCA without FA and MD resulted in a similar set of principal components accounting for 77.3% of variance and reinforcing our interpretation of the full model results. PC1_B_ accounted for 36.6% of variance and was analogous to PC2 from the full model, with loadings onto RD, NDI, and qihMT. PC2_B_ accounted for 22.7% of variance and loaded onto VF_m_ and g-ratio, similar to PC3. Finally, PC3_B_ accounted for 18.0% of variance and loaded onto AD and ODI, similar to PC1. Principal component analysis with varimax rotation is shown to be an effective way to collapse white matter imaging metrics into powerful, interpretable measures. Future studies employing this method should consider removal of broadly sensitive metrics such as FA and MD to improve specificity of resultant principal components.

Principal components were not significantly related to Total Reading in any investigated region. Bayes factors suggested the null hypothesis, no relationship between principal components and Total Reading, was substantially more likely than the alternative hypothesis in all regions. No relationships were identified in follow-up mixed effects models including principal components, age and scores from subtests included in the Total Reading composite score. Further, no correlations between initial measures and Total Reading were significant following multiple comparisons corrections. These findings suggest that gross relationships between white matter structural features and Total Reading ability are absent in typically developing adolescents, who tended to be skilled readers in our sample. Expansion of this analysis to a larger age range or comparison with a population with reading difficulty or dyslexia may provide a larger effect to assess, and is a promising direction for future multimodal investigations of white matter and reading.

Despite a lack of broad relationships between key white matter features and reading, some findings here hint that more specific relationships may be present in our sample. While no relationships were significant, p-values < 0.1 suggest a larger sample may find significant relationships between PC2 or PC3 and Total Reading in the left IFOF and left uncinate, respectively. Left hemisphere ventral white matter supports reading processing in skilled readers, and left inferior frontal regions have been consistently highlighted as related to reading skill in previous studies [3, 6, 8–10]. Additionally, qihMT was correlated with Total Reading ability in the bilateral arcuate fasciculus and ILF, the right IFOF and right uncinate fasciculus, and was trend level in the left IFOF. However, these findings did not survive multiple comparison corrections. Interestingly, qihMT was not significantly related to Total Reading in either the left IFOF or uncinate fasciculus, where trend level relationships with principal components were found. Trend level relationships between PC2, PC3, or qihMT and Total Reading provide some evidence for a link between axon diameter and myelin and adolescent reading. However, these relationships must be investigated and confirmed by future studies.

Links between principal components and age were identified throughout the brain. Relationships between PC2 and age were most prominent, found in all tracts except the uncinate fasciculus, and are visualized as scatterplots in Figure 4. Age-related trends tended to be similar between left and right hemispheres, suggesting that at the macro-scale, brain development is similar between hemispheres. This is in contrast to investigations of individual microstructural features, where increases in VF_m_ were shown to be largely left-lateralized during adolescence [57]. PC2 findings may be driven by NDI, as NDI has been previously shown to be age-sensitive and increases bilaterally throughout adolescence [57–59]. One relationship between PC1 and age remained in the left arcuate following multiple comparisons. While axon coherence tends to be stable across adolescence [60–62], we show that changes may still be ongoing in some regions. Gender was related to PC3 in the right inferior longitudinal fasciculus such that males had higher values than females. Higher PC3 values reflect higher VF_m_ and lower g-ratio values, thus the development of the right inferior longitudinal fasciculus may be further along in males. Studies of sex effects on white matter development have produced mixed results, suggesting either absence of or minor developmental effects during adolescence (for review see [63])., but large longitudinal studies remain necessary to effectively assess sex and gender effects across development.

## 5 CONCLUSIONS

Here, we have combined multimodal imaging techniques to assess white matter microstructure in reading-related white matter. Principal component analysis revealed three features of white matter microstructure which explained 79.5% of variance in our dataset. Principal components were related to tissue complexity, axon packing and myelin, and axon diameter, respectively. No significant relationships were observed between principal components and Total Reading, suggesting gross relationships between white matter structural features and reading are not present in typical adolescents. Some trend level results suggest minor roles for axon diameter and myelin in reading ability, but these findings must be confirmed by further research. Principal components are shown to be sensitive to age effects throughout the brain, and age findings were in line with previous studies applying PCA in white matter and other investigations of white matter microstructural development. Principal component analysis is an effective method to collapse multimodal sets of white matter imaging metrics into principal components which explain a large proportion of variance in white matter. Resultant principal components are age-sensitive and may prove useful to expand our understanding of links between white matter and reading in future studies. Use of such techniques to identify how white matter changes across the full developmental period, and how white matter structure is linked to various cognitive abilities, will provide an important baseline for future studies investigating developmental or cognitive disorders.

## ACKNOWLEDGEMENTS

This project was supported by the Natural Science and Engineering Research Council (NSERC). Salary funding was provided by NSERC (BG) and the Canadian Institutes of Health Research (CL). MC is supported by a Wellcome Trust Investigator Award (096646/Z/11/Z) to Derek K. Jones. The authors wish to thank the subjects and families who participated in this study, without whom this work would not have been possible.

